# Are the Next-Generation Pathogenicity Predictors Applicable to Cancer?

**DOI:** 10.1101/2024.05.06.592789

**Authors:** Daria Ostroverkhova, Yiru Sheng, Anna Panchenko

## Abstract

Next-generation pathogenicity predictors are designed to identify pathogenic mutations in genetic disorders but are increasingly used to detect driver mutations in cancer. Despite this, their suitability for cancer is not fully established. Here we have assessed the effectiveness of next-generation pathogenicity predictors when applied to cancer by using a comprehensive experimental benchmark of cancer driver and neutral mutations. Our findings indicate that state-of-the-art methods AlphaMissense and VARITY demonstrate commendable performance despite generally underperforming compared to cancer-specific methods. This is notable considering that these methods do not explicitly incorporate cancer-specific information in their training and have made concerted efforts to prevent data leakage from the human-curated training and test sets. Nevertheless, it should be mentioned that a significant limitation of using pathogenicity predictors for cancer arises from their inability to detect cancer potential driver mutations specific for a particular cancer type.

---

Many mutations contribute, or increase susceptibility, to various diseases, including cancer. Most of these mutations remain unannotated, and those that are characterized, tend to be disproportionately associated with certain protein groups^1^. It has been noticed that pathogenic missense mutations (resulting in changes of amino acids) are enriched at specific functional protein sites which has inspired the development of methods that try to decipher the functional effects of mutations on proteins. Various computational methods have been offered to rank mutations with respect to their pathogenic impacts^2^, and American College of Medical Genetics has recommended including several evidences from prediction tools to provide assessment of variants to be pathogenic or benign^3^.

The power of the new-generation pathogenicity predictors stands from the relatively large training data sets and the strength of machine learning models. These models are trained to learn high-order dependencies from protein structures and evolutionary constraints between amino acids using multiple sequence alignment of homologous proteins. In addition, the new generation pathogenicity prediction methods try to break the vicious circle of circularity and prevent the leakage of incorrectly annotated labels. They also leverage high statistical power by using the multitude of mutations observed in the human or primate population as benign and non-observed mutations as pathogenic.

Here we evaluate the two state-of-the-art representative methods of this group: VARITY^4^ and AlphaMissense^5^. VARITY specifically focuses on the annotation of rare variants and excludes any features derived from previous predictions. As to AlphaMissense model, it is pre-trained using AlphaFold architecture and uses labels assigned based on the minor allele frequency (MAF). Low frequency variants are treated as pathogenic, whereas benign variants are those having relatively high MAF in human and primate populations. VARITY and AlphaMissense utilize a diverse range of features such as residue co-evolution, structural constraints, evolutionary conservation, allele frequencies, among others.

Even though pathogenicity predictors primarily aim to predict pathogenic mutations in genetic diseases, these methods are routinely applied to detect somatic missense driver mutations in cancer patients. However, the logic of pathogenicity predictions is different from the cancer driver mutation detection. A pathogenic variant refers to a change in the DNA molecule that causes a malfunction of a protein leading to disease. In contrast, a cancer somatic driver mutation is a DNA alteration that actually confers a selective growth advantage to cancer somatic cells, thereby directly contributing to the development and progression of cancers. Furthermore, the underlying principle of classification of pathogenic variants differs from that for detecting cancer driver mutations. In the case of cancer driver mutation detection, the prediction framework is focused on distinguishing a few mutations which drive cancer development from a multitude of passenger mutations which do not lead to tumor growth in the same cancer patient and the same cancer type. On the other hand, pathogenic variants have a broader scope related to various diseases and are not necessarily cancer specific. These distinctions raise a question of whether the modern pathogenicity prediction methods can be effectively used in cancer studies to identify driver mutations.

Here we have evaluated the capability of AlphaMissense and VARITY for detecting cancer driver mutations. We then compared their performance to three representative non-ensemble methods that have been specifically designed to distinguish driver from passenger mutations in cancer: MutaGene^6,7^ FATHMM-cancer^8^, CHASMplus^9^. All these methods were chosen because they outperformed other approaches in recent studies^10,11^.

To compare and evaluate the performance of these methods, we used a comprehensive cancer experimental benchmark from the previous seven experimental studies^7,12^ with “driver” and “passenger” labels assigned following^7^. We also ensured that mutations from the benchmark were observed in the TCGA pan-cancer dataset^13^. As a result, the benchmark comprised 1048 missense variants with 613 positive (driver) and 435 negative (passenger) cases from 51 human genes (Table S1). We found that the performance of the five tested methods varied, with auROC ranging from 0.83 to 0.90 and Mathew’s correlation coefficient (MCC) varying from 0.55 to 0.68 (Figure 1A-B, Table S2). The highest performance was achieved by CHASMplus among all methods. For comparison, frequencies of observed mutations in cancer patients, often used as a prediction score in clinical settings, resulted in the lowest MCC score of 0.51 and auROC of 0.79 (Table S2).

**Figure 1.**
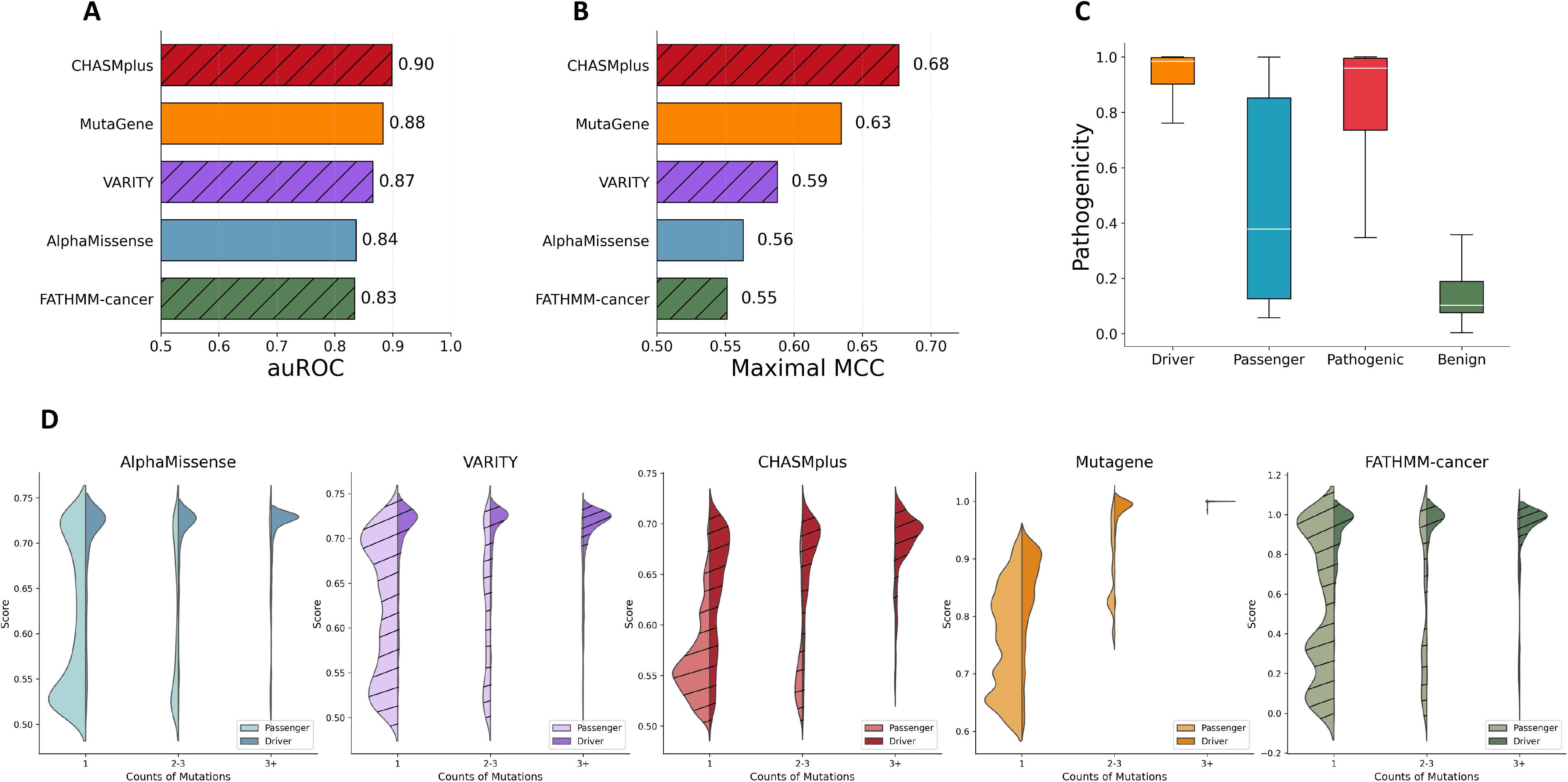
Comparative analysis of pathogenicity and driver mutation prediction methods. Assessment of performance of the computational methods at classifying cancer mutations using A) Area Under the Receiver Operating Characteristic Curve (auROC), and B) Maximum Matthew’s correlation coefficient (MCC). The bar marked with diagonal stripes indicates methods which can be compromised by data leakage between training and validation sets. The differences in auROC scores between VARITY and MutaGene, as well as between FATHMM-cancer and AlphaMissense, were not statistically significant based on the DeLong test. C) Distribution of the AlphaMissense pathogenicity scores for known driver (“Driver”), passenger (“Passenger”) mutations from the benchmark cancer dataset as well as for pathogenic (“Pathogenic”) and benign (“Benign”) mutations from the ClinVar dataset. The ClinVar dataset includes pathogenicity annotations for mutations related to various diseases and not limited to cancer. These annotations were obtained from either experimental validations or predictions. D) Distribution of the normalized prediction scores of five methods across three groups of mutations, each characterized by different mutation frequencies observed in cancer patients. The first group (“1”) contained 534 non-recurrent mutations, with 35% of known driver mutations among them, the second group (“2-3”) contained mutations recurring in two or three cancer samples with 71% of known drivers among them. Lastly, the third group (“3+”) had 94% of drivers out of mutations recurring in more than three cancer samples.

However, it should be mentioned that some results can be affected by the data leakage from the training sets. For example, approximately 19% of mutations in our benchmark were present in the ClinVar database^14^, which was utilized as the training set in VARITY. Furthermore, CHASMplus was trained on the TCGA mutation data (all mutations in our benchmark overlapped with the TCGA mutations) with various criteria applied for defining positive and negative class labels, whereas FATHMM-cancer model also trained its weights using data on cancer-associated mutations. This type of data leakage from training to the benchmark dataset may cause the overfit of the model and the corresponding methods are shown with diagonal stripes.

In general, the optimal performance of predictors is attained when the score distribution for positive cases is significantly shifted towards higher scores compared to negative cases. As observed in Figure 1C, using AlphaMissense as an example, the score distribution for benign ClinVar mutations was shifted to the lower scores and showed a minimal overlap with the distribution for pathogenic mutations. However, the score distribution for cancer passenger mutations was broader and partially overlapped with the tail end of the score distribution for driver mutations. This indicates that some cancer passenger mutations could potentially be misclassified by the method.

To further explore this issue, we have characterized mutations based on their occurrence frequency in cancer patients using the TCGA pan-cancer dataset (Figure 1D, Table S3). Our findings showed a consistent trend where recurrent mutations tended to have higher scores than non-recurrent ones across all five methods. Specifically, within the recurrent mutations class, driver mutations generally received higher scores than passenger mutations, as illustrated in Figure 1D. These observations suggest that all methods can effectively predict recurrent driver mutations. However, tools like AlphaMissense, VARITY, and FATHMM exhibited a second high-scoring peak within the score distributions for non-recurrent passenger mutations. This pattern indicates that these methods may be prone to falsely identifying some rare benign mutations as drivers.

Our findings indicate that the pathogenicity prediction methods, AlphaMissense and VARITY, demonstrate commendable performance despite generally underperforming compared to cancer-specific methods. This is notable considering that these methods do not explicitly incorporate cancer-specific information in their training. Developers of AlphaMissense, in particular, made a concerted effort to prevent data leakage from the human-curated training and test sets by labeling pathogenicity based on the MAF values.

Nevertheless, it should be mentioned that a significant limitation of using pathogenicity predictors for cancer arises from their inability to provide cancer type-specific predictions. This is critical because a mutation may act as a driver in one type of cancer but be a passenger in another. Overall, our evaluation underscores the potential for broader application of pathogenicity predictors in identifying cancer drivers, although this should be approached with caution.

## Materials and Methods

### Benchmark data set

To evaluate and compare the performance of five methods, we utilized a comprehensive benchmark dataset obtained from the seven experimental studies^12,15-20^ (https://github.com/Panchenko-Lab/Benchmark.git). The experimental evidence of impact of mutations included changes in enzymatic activity, response to ligand binding, impacts on downstream pathways, an ability to transform human or murine cells, tumor induction in vivo, or changes in the rates of progression-free or overall survival in pre-clinical models. The negative set mutations were those labeled as “neutral”, or without significant impacts on the normal protein functions or tumor formations.

### Mutation prediction methods

Five established computational methods were used for comprehensive evaluation of performance: MutaGene^6,7^, FATHMM-cancer^21^, CHASMplus^9^, VARITY^4^ and AlphaMissense^5^.

MutaGene uses the DNA context-dependent probabilistic models to estimate the nucleotide or codon level mutability to adjust the mutational observed recurrence to produce cancer type specific driver predictions. FATHMM-cancer algorithm is an adaptation of the original Functional Analysis through Hidden Markov Models (FATHMM) framework, which incorporates adjusted ‘pathogenicity weights’ for cancer-associated mutations. To calculate cancer-specific pathogenicity weights, FATHMM-cancer utilizes data on cancer-associated mutations from the CanProVar database^22^. The neutral polymorphisms are obtained from the UniProt database^23^.

CHASMplus utilizes a random forest algorithm to predict driver mutations in cancer. This method incorporates a variety of features derived from genomic and protein data, such as evolutionary conservation, structural impacts, molecular function annotations, and gene-level covariates. The TCGA mutation dataset was used to establish training labels. The positive class labels for missense mutations were selected based on three criteria: 1) occurrence in a curated list of 125 pan-cancer driver genes^24^, 2) occurrence in genes identified as significantly mutated in a specific cancer type according to MutSigCV^25^ and 3) occurrence in samples with relatively low mutation rate. The negative cases were the remaining missense mutations in the TCGA dataset.

The VARITY method employs a version of the gradient boosted tree algorithm (XGBoost). To train the VARITY model, a core and “add-on” training sets were assembled. The core training set consisted of high-quality missense variants, obtained from the ClinVar database and specifically limited to rare variants. Add-on training sets contained variants that might have less reliable information for determining their labels as positive or negative. The “add-on” training variants were obtained from various databases such as gnomDB^26^, HGMD^27^ and others. The VARITY model utilized a various set of features, but those informed by variant pathogenicity annotation or protein identity were excluded to limit the circularity.

AlphaMissense is known for the generation of a comprehensive database of predictions for all possible human single amino acid substitutions. AlphaMissense is based on a machine-learning model derived from the protein structure prediction capabilities of AlphaFold^28^. To mitigate the bias inherent in human-curated annotations, AlphaMissense employs weak labels derived from the population variant frequency data in humans and primates. Utilizing an unsupervised method to analyze population frequency data, AlphaMissense also identifies evolutionary patterns and functional characteristics of proteins. This strategy contrasts with the use of clinical data, which often introduces biases due to the uneven distribution of variant occurrences across genes.

### Statistical analyses and performance evaluation

The assessment of the prediction performance was done using auROC and the maximal Matthew’s correlation coefficient (MCC). The auROC evaluates the performance of a classification model, quantifying its ability to distinguish positive and negative cases correctly across all possible thresholds within each algorithm independently. The statistically significant difference between models was assessed using the DeLong test^29^ (Table S4), with a p-value of < 0.05 indicating a significant difference. The maximal Matthew’s correlation coefficient (MCC) provides a comprehensive measure of the quality of predictions, offering a balanced evaluation across various conditions.

## Acknowledgements

The authors were supported by the Department of Pathology and Molecular Medicine, Queen’s University, Canada. A.R.P. acknowledges the support of the New Frontier in Research Fund (NFRF) and the Natural Sciences and Engineering Research Council of Canada (NSERC). A.R.P. is the recipient of a Senior Canada Research Chair in Computational Biology and Biophysics. This study was conducted with the support of the Ontario Institute for Cancer Research through funding provided by the Government of Ontario. The views expressed in the publication are the views of the authors and do not necessarily reflect those of the Government of Ontario.

## Notes

### Competing Interest Statement

The authors have declared no competing interest.

